# CFTR corrector efficacy is associated with occupancy of distinct binding sites

**DOI:** 10.1101/2021.05.04.442442

**Authors:** Nesrine Baatallah, Ahmad Elbahnsi, Jean-Paul Mornon, Benoit Chevalier, Iwona Pranke, Nathalie Servel, Renaud Zelli, Jean-Luc Décout, Aleksander Edelman, Isabelle Sermet-Gaudelus, Isabelle Callebaut, Alexandre Hinzpeter

## Abstract

CFTR misfolding due to cystic fibrosis causing mutations can be corrected with small molecules designated as correctors. VX-809, an investigational corrector compound, is believed to bind CFTR directly to either the first membrane-spanning domain (MSD1) and/or the first nucleotide-binding domain (NBD1). Blind docking onto the 3D structures of these domains, followed by molecular dynamics (MD) simulations, revealed the presence of two potential VX-809 binding sites which, when mutated, abrogated corrector rescue. Mutations altering protein maturation are also shown to be not equally sensitive to the occupancy of the two sites by VX-809, with the most frequent mutation F508del requiring integrity of both sites and allosteric coupling with the F508del region while L206W only requires the integrity of the MSD1 site. A network of charged amino acids in the lasso Lh2 helix and the intracellular loops ICL1 and ICL4 is involved in the allostery between MSD1 and NBD1. Corrector VX-445, which is used in combination in clinics with VX-661, a structurally close analog of VX-809, to fully correct F508del, is also shown to occupy two potential binding sites on MSD1 and NBD1, the latter being shared with VX-809. In conclusion, VX-809 and VX-445 appear to bind different CFTR domains to alleviate specific folding defects. These results provide new insights into therapeutics understanding and may help the development of efficient corrector combinations.

## Introduction

Numerous mutations in the Cystic Fibrosis Transmembrane Regulator (*CFTR*) gene have been identified in patients affected with cystic fibrosis (CF). Mutations have been classified depending on the defect they cause, with the most severe forms of the disease being associated with premature termination codons (class I), protein misfolding (class II) or channel gating defects (class III). The most frequent mutation F508del causes multiple defects including protein misfolding, reduced channel activity and instability at the cell surface (*1*). Hence, the folding process of CFTR was extensively studied. Folding occurs co-translationally, including both intra- and inter-domain folding steps (*2, 3*). The first stages of co-translational folding leads to tight packing of the transmembrane (TM) helices of the first Membrane-Spanning Domain (MSD1) and assembly of its cytosolic N- and C-termini with the MSD1 first intracellular loop (ICL1), which is then strengthened by assembly with the first Nucleotide-Binding Domain (NBD1) (*4*). F508del, located in NBD1, was shown to alter both domain stability and inter-domain folding, mainly involving interactions with the ICL4 within the second MSD (MSD2) (*5*).

Early on, the folding defect associated with F508del was shown to be partially rescued by incubating the cells at 27°C (*6*), which is believed to stabilize key limiting folding intermediates and lead to the partial maturation of the protein. Rescue could also be achieved with small molecules such as chemical chaperones, one of the best examples being glycerol (*7*). These observations led to the screening of small molecules favoring the proper folding of mutant CFTR and to the identification of drugs named correctors, such as VX-809 (Lumacaftor) or its structurally closely related analogue VX-661 (Tezacaftor). While found very effective *in vitro*, their activity was suboptimal in clinical trials even in combination with the CFTR potentiator VX-770 (Ivacaftor) (*8, 9*), this latter enhancing channel function (*10*). To enhance rescue, corrector combinations targeting defects associated with different folding steps were identified by Veit and colleagues (*11*) showing synergistic rescue on CFTR-F508del. A corrector combination (VX-661 and a recently developed VX-445 (Elexacaftor)) in association with the potentiator VX-770 (triple combination therapy, called Trikafta™) showed enhanced rescue of CFTR-F508del and great clinical benefits of CF patients carrying at least one F508del allele (*12*).

VX-809 and VX-661 are believed to target the same CFTR region. Nonetheless, as these molecules have been identified using high throughput screening, their mechanisms of action and binding sites remain largely elusive, as for the recently described VX-445 (*12, 13*).

VX-809 was shown to stabilize truncated CFTR fragments that only contain the entire N-terminal MSD1 domain (*4, 14, 15*). Corrective conformational changes or stabilization of MSD1 by VX-809 appears then to be communicated to other domains, including NBD1, MSD2 and NDB2. VX-809 is otherwise highly effective at correcting MSD1-specific disease-causing mutations (*16, 17*), some of them forming a hot spot around a groove predicted on the 3D structure of the protein to bind phospholipids in a stable way (*18*). These mutations include P67L and L206W (*11, 15, 17, 19–21*) with the notable exception of G85E which was shown to be resistant to correction (*4, 22*). This groove, which can therefore be considered as the MSD1 Achilles heel, is a potential binding site for small molecules that could act by stabilizing the whole MSD1 domain during the folding process. It differs from the composite, multi-domain binding pocket proposed by Molinski and colleagues (*23*) encompassing amino acids F374-L375, and amino acids from the lasso, NBD1, MSD2 and ICL4.

Here, we show that VX-809 can be docked and is stably accommodated in this MSD1 groove and that mutations of some critical amino acids modify the sensitivity to VX-809 correction. We also describe an allosteric coupling linking this site to the F508del region on NBD1. In addition we highlighted a second potential VX-809 binding site within NBD1, whose existence is supported by previous experimental evidence of a direct binding of the drug to this domain and allosteric coupling to the NBD1:ICL4 interface which is directly affected by the F508del mutation (*24*). While MSD1 mutants prevented both F508del and L206W rescue by VX-809, NBD1 mutants prevented rescue of only F508del. These results are consistent with the presence of two VX-809 binding sites and reveal that class II mutants present a different sensitivity to corrector single or double occupancy. The sensitivity of these mutants to the novel corrector VX-445 and VX-809/VX-445 combination was also consistent with the presence of two binding sites also located on MSD1 and NBD1 and the need of an allosteric coupling between the two sites to fully correct F508del.

## Results

### A potential VX-809 binding site highlighted within MSD1 by molecular docking

We have previously shown that cis mutations of F508del could alter the response to VX-809 treatment, either by inducing an additional folding defect as for L467F (*25*) or by altering a potential VX-809 binding site as suggested with F87L (*26*). Here, we evaluated the effect of L53V, another cis mutation located in the N-terminal part of CFTR, called the lasso motif and defined after the first high-resolution experimental 3D structures of zebrafish and human CFTR (*27, 28*). In transiently transfected HEK293 cells, CFTR-L53V showed a maturation profile comparable to WT with the presence of both fully-glycosylated band C and partially glycosylated band B at a similar ratio (Fig. 1 (A and B)). In combination with F508del, L53V prevented VX-809 rescue, similarly to F87L (Fig. 1 (A and B)). These results led us to further evaluate the region in the vicinity of amino acids L53 and F87-for the presence of a potential VX-809 binding site.

**Figure 1.**
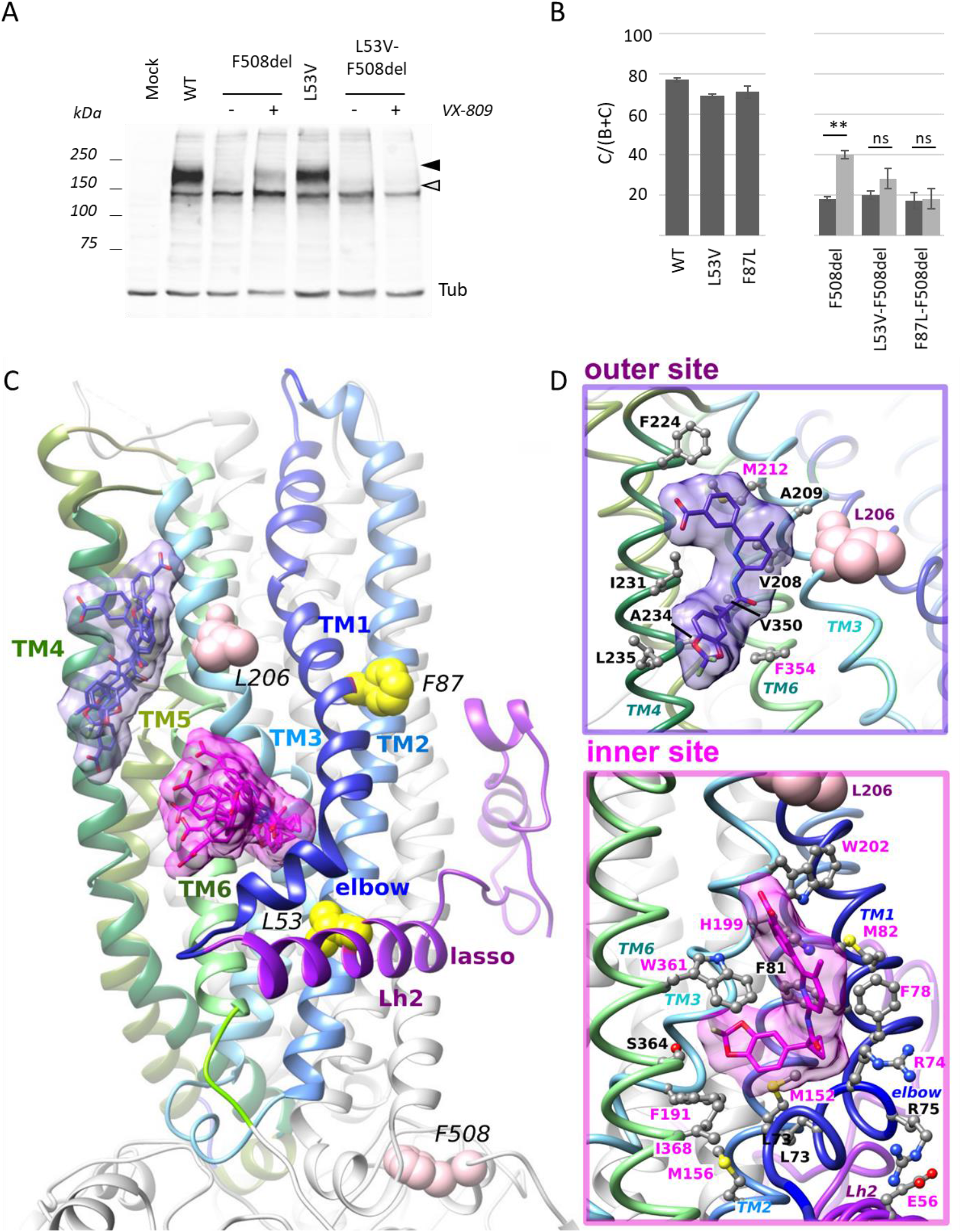
Identification of a potential VX-809 binding site with MSD1. **A**. Western blot analysis of WT and mutant CFTR. Fully glycosylated and core glycosylated CFTR are indicated with a dark grey and light grey arrow. Cells were treated with VX-809 (3μM, 24h) when indicated. Tubulin (Tub) was probed to assess equal loading amounts. **B.** Quantification of the C/(B+C) maturation ratio of the indicated mutants. Dark grey represents control conditions and light grey VX-809 treatments. Measures are means ± SEM of n=3-20, with * indicating p<0.05, ** indicating p<0.01 and ns indicating no statistical differences (one-way ANOVA followed by Fischer test for p evaluation). **C.** Two main pockets were identified at the level of the membrane inner and outer leaflets on the 3D structure of human CFTR MSD1, extracted from the cryo-EM 3D structure of the full-length protein (pdb: 6MSM). The membrane inner leaflet site corresponds to the best docking scores obtained after blind docking using SwissDock, with the surface envelope of VX-809 (representative members of the 6 first clusters) shaded in pink. The second binding site was identified at the level of the membrane outer leaflet, with fewer clusters (surface envelope (representative members of 3 clusters) shaded in purple) and less favorable docking scores. **D.** Focus on amino acids within the potential VX-809 binding sites at the level of the membrane inner (bottom) and outer (top) leaflets. The conformations of VX-809 corresponding to the best docking scores in the two sites are shown (inner site (cluster 0): *SwissDock FullFitness* score of −1938; outer site (cluster 7): *SwissDock FullFitness* score of −1924). Tested amino acids are labeled in pink.

Blind docking of VX-809 was performed on the experimental 3D structure of the human ATP-bound CFTR (PDB 6MSM (*29*)) using the EADock DSS tool provided by the online SwissDock server (*30, 31*). Only the MSD1 was considered as a target for docking with default parameters. Clusters gathering similar conformations (or binding modes) of VX-809 within sites were ranked according to the average FullFitness of their elements (*32*). After having excluded one unauthorized site in the full-length CFTR, only two binding pockets were identified on the MSD1 surface, totalizing 17 and 11 clusters respectively (pink and purple surfaces in Fig 1C). The first pocket, fetching the first six best clusters, is located at the level of the membrane inner leaflet and formed by amino acids from the elbow (L73), the N-terminal parts of TM1 (F78, F81, M82) and TM3 (F191, H199, W202) and the C-terminal parts of TM2 (M152, M156) and of TM6 (W361, S364, I368) (Fig. 1D-bottom, Fig. S1A). All the binding modes for these six first clusters, including a total of 43 conformations, are characterized by a unique position of the benzodioxolyl core of VX-809, deeply embedded in the pocket, as it is also the case for the amide group, while the 2-phenylpyridyl portion and the carboxylate group adopt more variable positions, outside the pocket (Fig. S1B-left). Worth noting is that a similar blind docking of VX-661 on MSD1 led to highlight the same binding pocket in the first five best clusters, with a very similar position of the common benzodioxolyl core, whereas variable positions are observed outside the pocket for the second half of the drug (Fig. S1B-right). Hence the difluoro benzodioxolyl group is the key anchor element of the two correctors in the binding pocket. This site is embedded within a hot spot for class II mutations associated with either mild folding defects (in pink on Fig. S2: P67L, E56K, V232D, I336K, Q359K, T360K, R347P, S341P and I336K, and W361R) or severe folding defects (in red: G85E, E92K, G91R, H199Y, P205S, L206W and L227R) (*7, 18*), consistent with a fragility in the folding and/or packing of the transmembrane helices (TM) in this region.

The second site, highlighted at the level of the membrane outer leaflet, involves amino acids of the C-terminal part of TM3 (V208, A209, M212) and N-terminal parts of TM4 (F224, I231, A234, L235) and TM6 (V350, F354) (Fig. 1D - top). This region appears less fragile as only two variants of unknown significance have been referenced in the Sickkids database (http://www.genet.sickkids.on.ca/), namely A209S and A234D.

### Exploring key residues of the MSD1 VX-809 binding pockets

We further explored the two potential pockets by site-directed mutagenesis of key amino acids implicated in VX-809 binding and evaluated their impact on correction on two class II mutants, F508del which is partially rescued and L206W which can be fully rescued (Fig. 2 (A and B)). The potential binding site located at the level of the membrane outer leaflet (Fig. 1D) was evaluated by generating mutants M212A and F354A, opposite of each other. These mutations did not alter protein maturation by themselves (Fig. S3A) nor prevent VX-809 correction of both F508del (Fig. 2A) and L206W (Fig. 2B), indicating that mutations within this site do not preclude VX-809 induced correction.

**Figure 2:**
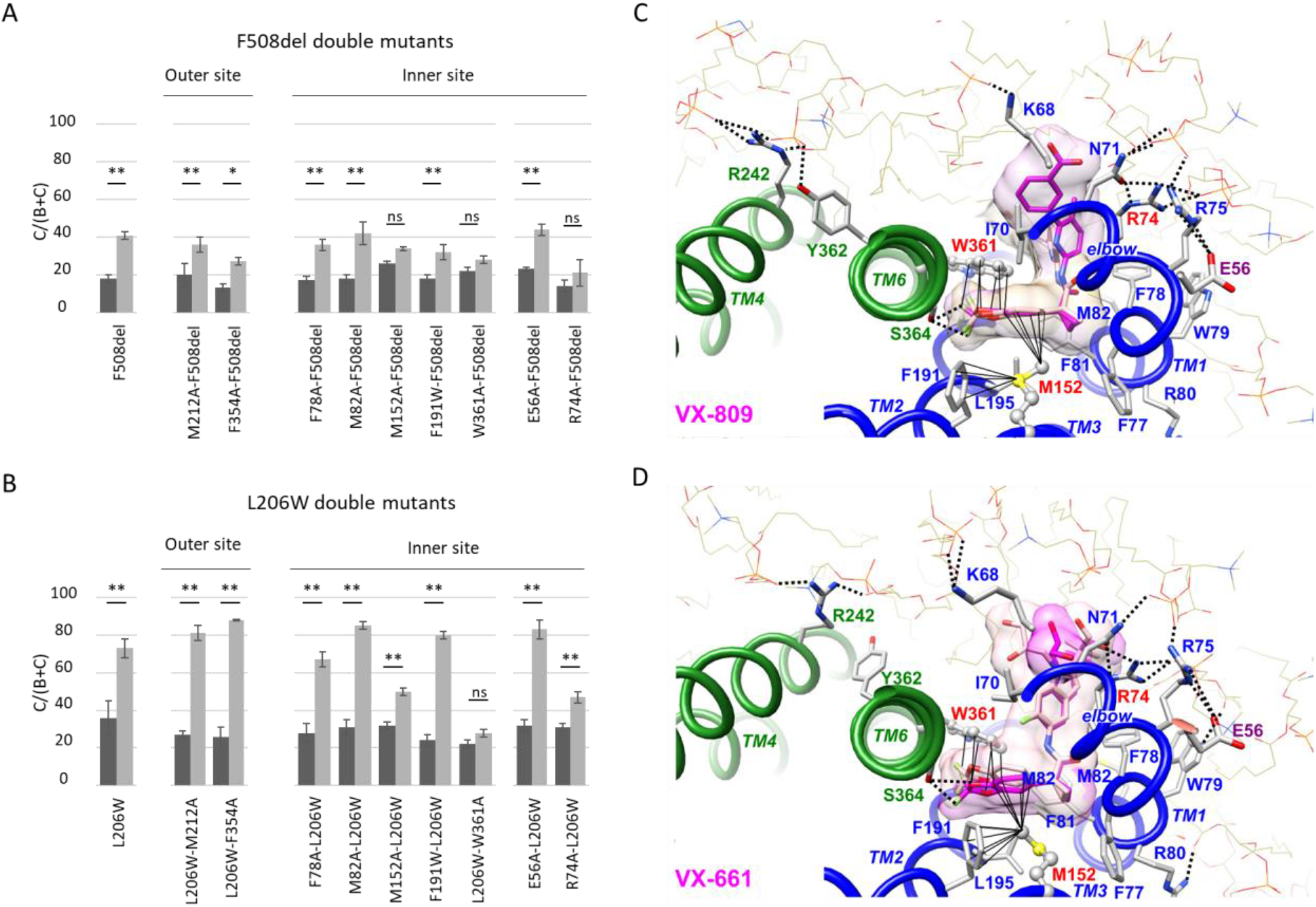
Exploring MSD1 inner membrane leaflet binding site. **A-B.** Quantification of the C/(B+C) maturation ratio of the indicated mutants. Dark grey represents control conditions and light grey VX-809 treatments. Measures are means ± SEM of n=3-20, with * indicating p<0.05, ** indicating p<0.01 and ns indicating no statistical differences (one-way ANOVA followed by Fischer test for p evaluation). **C.** Focus on the VX-809 position within the membrane inner leaflet binding site, before (light pink) and after (dark pink) MD simulation (125 ns), highlighting the stable position of the drug in the binding site and interactions with lipids (POPC) covering the site. The tight contacts established with M152 and W361 are also represented with black lines. Two H-bonds are present between S364 OG and VX-809 F1 (3.08 Å) and F2 (3.02Å). **D.** Similar view for VX-661, before (light pink) and after (dark pink) MD simulation (125 ns). Two H-bonds are present between S364 OG and VX-661 F3 (2.98 Å) and O3 (3.02Å).

The pocket located at the level of the membrane inner leaflet was evaluated by mutating a set of nine amino acids (F78A, M82A, M152W/A, M156W, F191W, H199W/A, W202A, W361A and I368W). Six out of the 11 tested mutations induced significantly decreased maturation by themselves (M152W, M156W, H199W, H199A, W202A and I368W, Fig. S3B), confirming the vulnerability of this region. Single mutants M156W, H199A and I368W could be rescued by VX-809 up to WT levels while M152W, H199W and W202A showed only a partial or no rescue, consistent with these amino acids being implicated in VX-809 induced correction (Fig. S3C).

The effect of F78A, M82A, M152A, F191W and W361A that were not deleterious in the WT context was then tested on the rescue of F508del (Fig. 2A) and L206W (Fig. 2B). While F78A, M82A and F191W had no effect, secondary mutations M152A and W361A reduced or prevented CFTR rescue by VX-809. The importance of M152 and W361 in VX-809 binding was further strengthened by their position on both sides of the difluoro benzodioxolyl group, which is the key anchor point of the drug in the site (Fig. 1D). Molecular dynamics (MD) simulation indicated a very stable position of VX-809 within the binding site, with H-bonds between the fluor atoms and S364 hydroxyl, as well as a particular involvement of lipids covering the 2-phenylpyridyl portion of the drug and establishing, through their phosphate groups, key interactions with several basic amino acids (Fig. 2C). A similar situation is observed for the MD simulation of the CFTR:VX-661 complex (Fig. 2D). To assess if M152A and W361A altered VX-809 binding affinity, higher VX-809 concentrations were used. While CFTR-M152A-F508del was partially rescued at high concentrations, CFTR-W361A-F508del remained not responsive (Fig. S4 (A, B and C)). For secondary mutations which induced CFTR misfolding (M152W, M156W, H199W/A and I368W), both F508del and L206W correction was prevented (Fig. S3 (D and E)), in this case probably due to additive folding defects, as was previously demonstrated for cis mutation L467F (*25*). Of note, cis-mutations F87L and L53V which prevented VX-809 correction of F508del CFTR are located in the vicinity of the inner binding site (Fig. 1C), and may act on this one by allostery. The F87 side chain is indeed interacting with that of F83, two amino acids downstream F81, whereas L53 is in contact with M156 and C76 (Fig. 1D-bottom).

We then targeted amino acids not lying directly within the membrane inner leaflet pocket but implicated in its architecture. Amino acids E56 and R74 are located at the cytoplasmic side of the pocket, either at its edge (R74) or at the interface between the elbow and the lasso (E56). E56 forms a salt-bridge with R75 (Fig. 1D). We generated E56A and R74A in WT, F508del and L206W plasmids. Single mutation E56A and R74A did not alter protein maturation (Fig. S3B). While E56A had no apparent effect, R74A prevented F508del correction (Fig. 2A) and mitigated L206W correction (Figs 2B). Treatment of CFTR-F74A-F508del with higher VX-809 concentrations did not rescue maturation (Fig. S4D).

Taken together, these results confirm the sensitivity of this region to mutations and show that amino acids directly located in the membrane inner leaflet pocket of MSD1 (M152, W361) or involved in its architecture, especially by establishing bonds with lipids (R74), influence VX-809 efficacy towards the two class II mutations, supporting thereby its relevance as a VX-809 binding site.

### Allosteric coupling with F508del

In order to understand the impact of a binding site in MSD1 on the NBD1:ICL4 interface affected by the F508del mutation, we searched for a possible allosteric coupling between the MSD1 VX-809 binding site and NBD1 (Fig. 3A). A network of connected amino acids could be highlighted, including a salt-bridge formed between amino acids K162 (in ICL1, at the end of TM2) and E1075 (in ICL4, at the beginning of TM11), the latter being connected to the NBD1 F508 region (Fig. 3A). This salt bridge, not present on the cryo-EM 3D structure (Fig. S5A-top), appeared quickly in the 1μs long MD simulation run on the experimental 3D structure of CFTR and remained stable over time (Fig. S5A-middle). It adds up to the salt-bridges linking K163/K166 (ICL1) to E379 (at the end of TM6) (Fig. S5A-top and middle), thereby reinforcing the network linking MSD1 to the ICLs. After MD simulation in presence of VX-809, the K162-E1075 salt-bridge remained very stable as well (Fig. 3A and Fig. S5A-bottom). Remarkably, the occupancy of the MSD1 inner leaflet binding site by VX-809 is accompanied by a movement of the lasso Lh2 relative to ICL1-ICL4, leading K162 to shift an additional H-bond from E54 to D47. Analysis of the 1μs long MD simulation run on CFTR in absence of VX-809 clearly indicated a community of highly correlated residues in the vicinity of the MSD1 VX-809 binding site and ICL4 (in contact with NBD1 F508) via the lasso and ICL1, that is disrupted by the double mutant K162A-E1075A (Fig. S5B). Altogether, these results strengthen the importance of the lasso Lh2 for allosteric coupling between MSD1 and NBD1, displaying a tight network of salt-bridges involving the potential MSD1 VX-809 binding site (through the E56-R75 bond) and ICLs (through the E54/D47-K162-E1075 network), which leads in turn to impact the ICL4:NBD1 interface involving F1074 (Fig. 3A and Fig S5A). The VX-809 binding site is also connected to ICL1 through M152 (on the same TM2 helix than K162) and through the bond network linking the end of TM6 to K163/K166. Maturation of mutant K162A was reduced while E1075A was not altered (Fig. 3C). Both K162A and E1075A prevented F508del rescue while not affecting L206W correction (Fig. 3 (D and E)). Treatment of CFTR-F508del-E1075A with higher VX-809 concentrations did not rescue maturation (Fig. S4E) while swapping amino acids K162E-E1075K of the salt bridge restored VX-809 rescue (Fig. 3D). These results are consistent with the necessity of an allosteric coupling between MSD1 and mutant NBD1 to rescue F508del, contrary to L206W. Allosteric coupling towards NBD1 could also explain the differential effect of cis-mutations L53V and F87L in combination with either F508del (Fig. 1B) and L206W (Fig. 3D).

**Figure 3.**
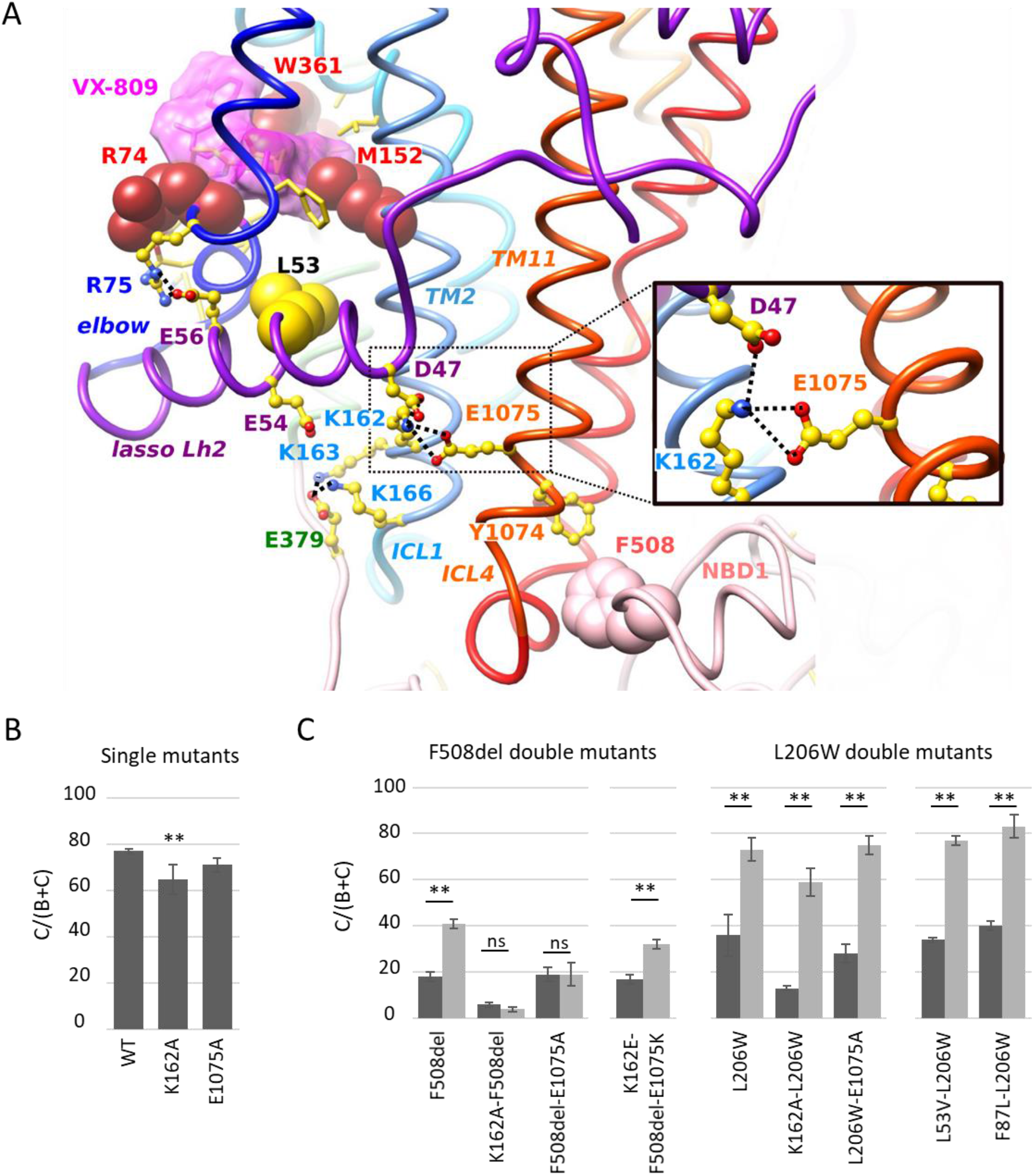
MSD1 binding site and F508 region are allosterically coupled. **A.** Path linking the potential VX-809 binding site in MSD1 and the F508 position at the interface between NBD1 and ICL4 and involving lasso Lh2 and ICL1 (3D structure after MD simulation in the presence of VX-809). The surface envelope of VX-809 is pink shaded. The K162-E1075 bond network is highlighted in the inset. **B-C.** Quantification of the C/(B+C) maturation ratio of the indicated mutants. Dark grey represents control conditions and light grey VX-809 treatments. Measures are means ± SEM of n=3-20, with * indicating p<0.05, ** indicating p<0.01 and ns indicating no statistical differences (one-way ANOVA followed by Fischer test for p evaluation).

### A second potential VX-809 binding site on NBD1

As a VX-809 binding site was also highlighted on NBD1 (*24*), we performed blind docking of VX-809 on the crystal structure of human F508del CFTR NBD1 (pdb:4WZ6), from which the regulatory extension has been removed. The regulatory extension does not belong to the NBD1 domain and has a highly flexible position, being located in the crystal structures of isolated NBD1 at the level of the NBD1:NBD2 interface. Isolated NBD1 3D structure however has a definite position for the last short α-helix from NBD1 (H9: aa 638 to 645), which has not been captured in the cryoEM 3D structure. While the two first docking binding modes are located at the level of the F508 loop (cluster 0: blue in Fig. 4A) and ATP-binding site (cluster 1: green), respectively, the following 7 clusters (orange), gathering a total of 65 conformations, focused on a single region, involving strands S3, S9, S10 and helices H3, H8 and H9. Representative conformations of each of the 7 clusters are given in Fig. 4A. The binding site, similar to the one proposed by Hudson and colleagues (*24*) and including amino acid H620, is oriented towards the interface that NBD1 makes with NBD2 and is located in a region also highlighted previously as critical for CFTR trafficking (*33*). MD simulation of the VX-809:NDB1 complex, considering the best docking score (cluster 2), led to refine the position of the drug in the binding site, and highlight the critical role of H620 and Y625, as well as basic amino acids of which K643 (Fig. 4A). A strong H-bond is observed between VX-809 O5 and Y625 hydroxyl group, while significant interactions also exist between the two VX-809 fluor atoms and S623 hydroxyl group. Of note is that a similar docking of VX-661 led to highlight the same binding pocket in six of the ten best clusters (the four first ones corresponding to the F508 loop), supporting the relevance of this site for corrector targeting (Fig. S6).

**Figure 4.**
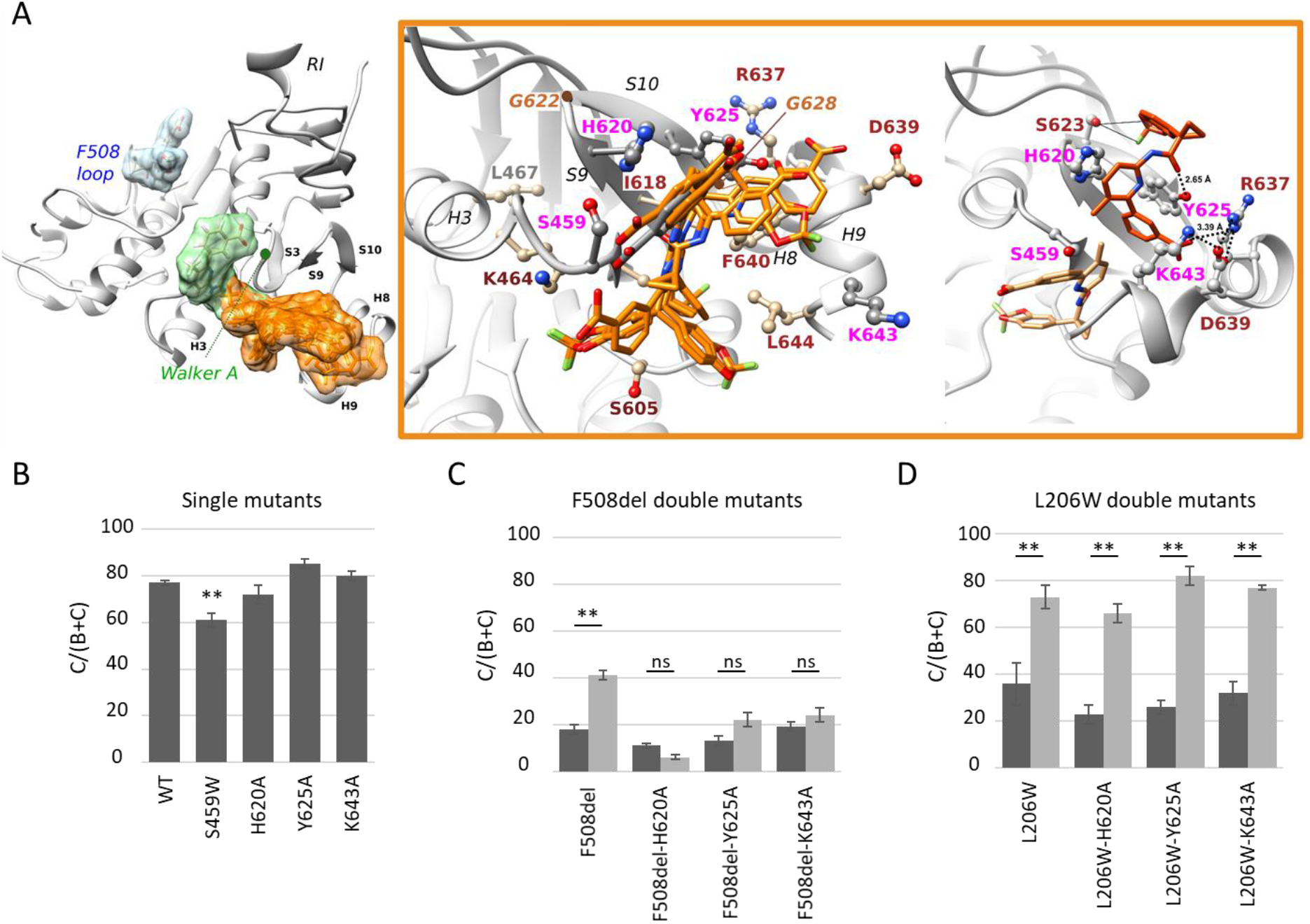
An intact NBD1 binding site is required for F508 rescue. **A. Left:** View of the human CFTR F508del NBD1, extracted from a crystal structure (pdb: 4WZ6) from which the regulatory extension (RE) has been removed and on which VX-809 has been docked (blind docking). Three main pockets were identified in the high-ranking clusters, the third one (orange) gathering 7 clusters out of 9 (*SwissDock FullFitness scores of −1274 to −1270*). **Middle:** Representative conformations of each of the 7 clusters of the third pocket, with its main amino acids highlighted in ball and stick. The positions of G622 and G628, identified as critical for CFTR trafficking (33), are indicated with brown circles. **Right:** Position of VX-809 (cluster 2) in the NBD1 binding site, before (light pink) and after (dark pink) MD simulation, highlighting the refined position of the drug and tight contacts established with H620 and Y625, as well as with S623 and basic amino acids belonging to H8 and H9. **B-C-D.** Quantification of the C/(B+C) maturation ratio of the indicated mutants. Dark grey represents control conditions and light grey VX-809 treatments. Measures are means ± SEM of n=3-20, with * indicating p<0.05, ** indicating p<0.01 and ns indicating no statistical differences (one-way ANOVA followed by Fischer test for p evaluation).

A set of mutations was introduced within this potential NBD1-located VX-809 binding site (labeled pink in Fig. 4A). H620A, Y625A, and K643A did not alter CFTR processing while S459W reduced maturation without preventing VX-809 rescue (Figs. 4B and S2E). Strikingly, VX-809 rescue of F508del and L206W were differently affected by these mutations. Indeed, while S459W, H620A, Y625A and K643A prevented F508del correction, they did not affect L206W restoration (Figs 4 (C and D) and S3F). Treatment of CFTR-F508del-H620A with high VX-809 concentrations did not rescue maturation (Fig. S4E). These results indicate again that the two class II mutants are not corrected by the same mechanism and suggest that an intact NBD1 binding site is necessary for F508del rescue, contrary to L206W. These differences could be explained by the distinct folding defects associated with F508del and L206W. Contrary to L206W, F508del was shown to destabilize NBD1 (*34*).

### Some of the highlighted sites can also accommodate corrector VX-445

CFTR-F508del correction is greatly enhanced by the combination of two correctors, *e.g*. VX-809/VX-445 or VX-661/VX-445 with the later showing clear clinical benefits (*12*). We evaluated the sensitivity of our mutants to corrector VX-445 and to the VX-445/VX-809 combination. VX-445 enhanced CFTR-F508del maturation alone, reaching comparable rescue to VX-809 and up to near WT levels when in combination with VX-809 (Fig. 5A). Cis mutations identified in CF patients affected differently treatment response. While L53V and F87L did not affect correction by VX-445 and VX-809/VX-445, L467F prevented correction in both conditions (Fig. S7).

**Figure 5.**
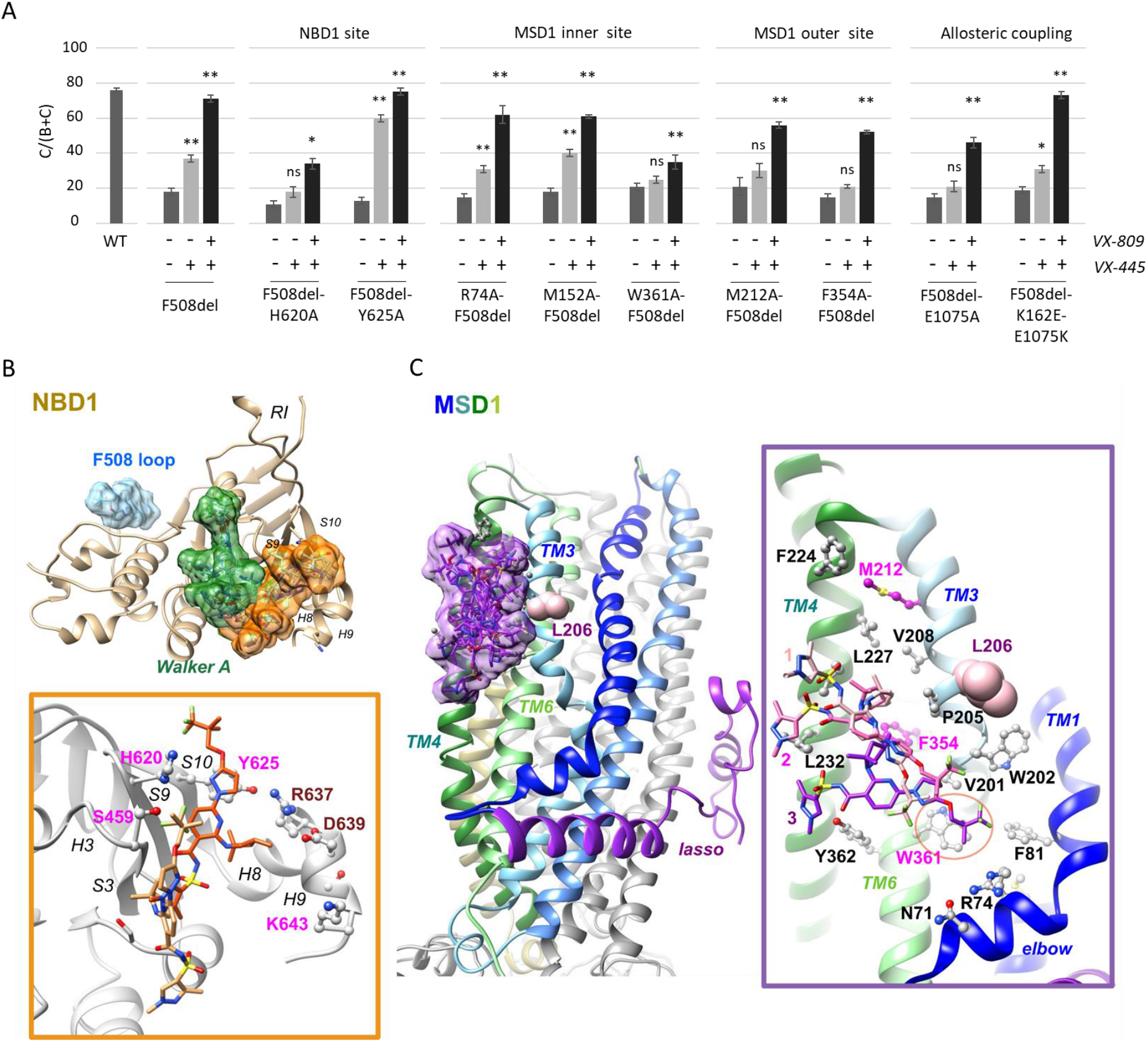
Rescue by VX-445 and VX-809/VX-445 requires both MSD1 NBD1 integrity. **A.** Quantification of the C/(B+C) maturation ratio of the indicated mutants. Dark grey represents control conditions and light grey VX-809 treatments. Measures are means ± SEM of n=3-20, with * indicating p<0.05, ** indicating p<0.01 and ns indicating no statistical differences (one-way ANOVA followed by Fischer test for p evaluation). **B-C.** Blind docking of VX-445 on NBD1 (B) and on MSD1 (C), followed by MD simulations. For NBD1, three main pockets were identified in the ten high-ranking clusters, as for VX-809, the third one (orange) gathering 5 clusters out of 10 (*SwissDock FullFitness* scores of −1662 to −1656), whereas for MSD1, only one allowed pocket was highlighted (first 8 clusters shown, *SwissDock FullFitness* scores of −2324 to −2313). Representative conformations of the first clusters in MSD1 and in NBD1 (orange site) are given in Fig. S8. In the boxes are shown the positions of VX-445 in the NBD1 binding site (cluster 2) and in the MSD1 binding site (cluster 8), before (light colors) and after (dark colors) MD simulation (125 ns), highlighting the refined positions of the drug within the sites. An intermediate position of VX-445 after 60 ns (2), between 0 ns (1) and 125 ns (3) was indicated for the MSD1 binding site. The orange circle in the MSD1 box highlights interaction of W361 with the 3,3,3-trifluoro-2,2-dimethylpropoxyl group of VX-445.

For secondary mutations located within NBD1, Y625A did not affect treatment responses while H620A prevented VX-445 and blunted VX-809/VX-445 combination correction (Fig. 5A). This led us to suppose that VX-445 may occupy a binding site in the vicinity of H620. This hypothesis was supported by blind-docking of VX-445 on NDB1, highlighting a preferential binding site similar to that targeted by VX-809 (Fig. 5B). As for VX-809, molecular dynamics simulation of the NBD1 with VX-445 (best docking score), led to refine the position of the drug in the binding site in a similar way, and also highlighted the critical role of H620 in drug contact (Fig. 5B).

In the context of MSD1 secondary mutations R74A and M152A, both VX-445 and VX-809/VX-445 corrections were maintained while W361A was found to abrogate VX-445 correction and markedly reduce VX-809/VX-445 correction (Fig. 5A). These results suggested that W361A may affect a binding site within MSD1 possibly distinct from the VX-809 site embedded in the inner leaflet as neither R74A nor M152A altered correction. Blind-docking of VX-445 on MSD1 identified a unique potential VX-445 binding site (Fig. 5C), corresponding to the site at the outer membrane leaflet described in Fig. 1C for VX-809. While substitutions M212A and F354A located within this site did not prevent VX-809 correction (Fig. 2A), they abolished VX-445 correction and reduced the level of correction obtained with the VX-809/VX-445 combination (Fig. 5A).

MD simulations of CFTR withVX-445 bound to this outer leaflet site showed that one binding pose, for which the 3,3,3-trifluoro-2,2-dimethylpropoxyl extremity could be potentially in contact with the other inner leaflet binding site, evolved towards a stable position after ~100 ns (illustrated here at 125 ns, as for all the MD simulations of protein-drug complexes, Fig. 5C). This position implied a shift towards W361, with which VX-445 tightly interacts, however without reaching the VX-809 binding site. MD simulation considering another pose, orientated along the same axis but in an opposite way, led to a rapid escape of the drug out of the binding site. We also ran MD simulations with correctors in both sites (*i.e*. VX-445 in the outer membrane leaflet site and VX-809 or VX-661 in the inner membrane leaflet site) and observed in this case a stable binding of VX-809/VX-661, as previously observed, together with a stable position of VX-445 (Figs 6 and S9). Correctors interacted with each other through their extremities, consistent with a synergic behavior. The VX-661/VX-445 couple showed a qualitatively better complementary to the protein surface, with strong interaction between VX-661 and R74 (Fig. S9). Worth noting is that E1075A which interrupted the allosteric coupling between MSD1 and NBD1 (Fig. 3 and S3), also prevented VX-445 and VX-445/VX-809 correction while reconstitution of the K162-E1075 salt bridge (K162E-E1075K) restored full correction (Fig. 5A). This result highlights the importance of the K162-E1075 salt bridge for F508del correction by both VX-809 or VX-661 in association with VX-445 (Fig 6). As for VX-809 and VX-661 alone, slight local conformational changes were observed (Fig. 3 and S5), in particular in the elbow helix and the lasso Lh2, allowing D47 to bind K162 in ICL1, which forms a salt bridge with E1075 in ICL4.

**Figure 6:**
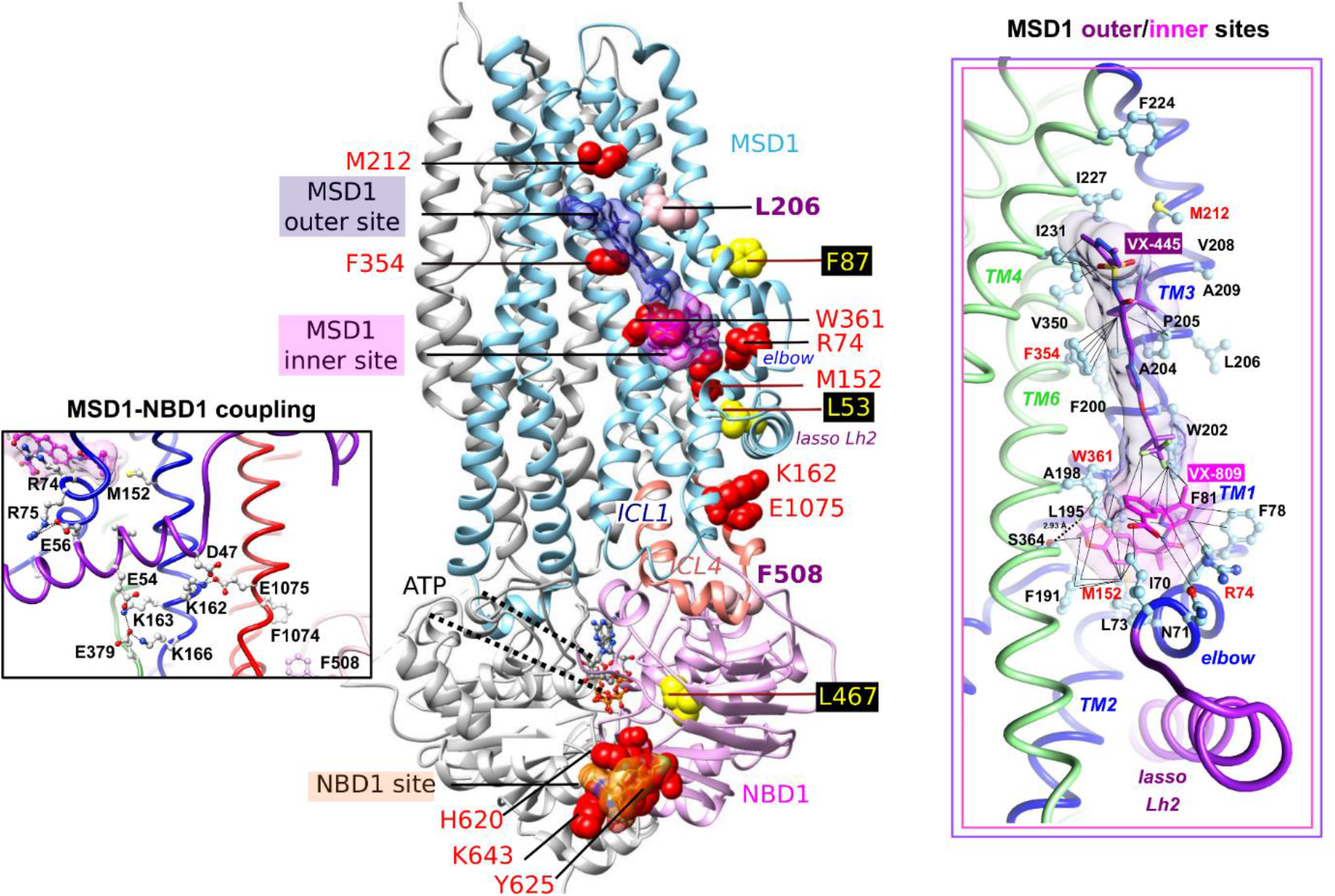
Potential corrector binding sites on the CFTR 3D structure. View of the 3D structure of human CFTR (ribbon representation, pdb 6MSM), on which the correctors are highlighted in their stable positions (after 125 ns MD simulations) in their potential binding sites using grids (pink and orange for VX-809/VX-661 and purple for VX-445). Amino acids L206 and F508 are indicated in pink. Cis mutations identified in the context of complex alleles which prevent VX-809 rescue of CFTR-F508del are represented in yellow. Secondary mutations preventing the correction of CFTR-F508del or both CFTR-L206W and CFTR-F508del are in red. The 3D structure of human F508del NBD1 (pdb:4WZ6) has been considered for NBD1 helices H8/H9, including the K643 position. At right are shown with atomic details the positions and contacts of VX-809 and VX-445, considered simultaneously, into the inner and outer membrane sites, respectively (also see Fig. S9 for detailed comments and comparison with the VX-661/VX-445 complex). At left is represented the bond network involved in the coupling between MSD1 and NBD1 (illustrated in the CFTR-VX-661/VX-445 complex, also see Fig. S10 for a view of the network in the different complexes).

Taken together, these results suggest the presence of two binding sites for each of the correctors, located on MSD1 and NBD1.

## Discussion

Several small molecules targeting CFTR have been identified in the past decade, enabling to enhance channel activity, protein stability or folding. These molecules likely bind different sites on the CFTR protein and have various effects on CFTR mutations, supporting “theratyping” of rare alleles according to pharmacological responses (*1, 7*) and the need of a better understanding of the molecular mechanisms involved in therapeutical strategies.

Corrector VX-809 has been developed first as it partially restores trafficking of F508del CFTR (*35*). VX-809 does not improve isolated NBD1 stability (*14*) and evidence have been provided that it may act by stabilizing the NBD1:ICL4 interface (*2, 3, 16*). Some studies have suggested a possible binding site at the level of this interface (*e.g*. (*5, 36*)), which is supported here by the fact that the best scoring pose of VX-809 on the F508del NBD1 3D structure indeed fills this interstitial area (Fig. 4A), as also proposed for the c407 corrector (*37*). However, this seems specific to F508del CFTR, as VX-809 was also shown to be efficient on a wide range of class II mutations which are located on other CFTR domains (*11, 15, 38, 39*), while the NBD1:ICL4 interface is intact. The various locations of the VX-809 rescuable mutations on the whole CFTR protein, in particular in MSD1, is in favor of allosteric effects and/or multiple binding sites for this small drug.

Two seminal works have shown that VX-809 bind to both MSD1 (*15*) and NBD1 (*24*), inducing different effects. Binding to MSD1 was suggested by the stabilization of truncated CFTR forms, encompassing amino acids 1-373, likely by favoring a folding intermediate (*15*). This stabilizing effect appears to be sufficient to achieve a strong rescue of class II mutants located in the N-terminal part of CFTR, such as E56K, P67L, L69H, L206W or W361R (*11, 15, 18–20*), indicating that targeting initial MSD1 folding and assembly accounted for the bulk VX-809 rescue. The VX-809-binding site on MSD1 has remained heretofore elusive. Molinski and colleagues have proposed a potential binding site located close to the cytoplasm and involving interfaces with other domains (*23*). Nonetheless, mutants in this site (K166E/Q) showed strong response to VX-809 (*4*). We followed here a strategy consisting in using MSD1 cis mutations identified in patients carrying F508del to identify amino acids modulating the VX-809 response (Fig. 1 and Fig. 6). Both L53V and F87L prevented F508del rescue and are located in the vicinity of a potential binding pocket, which was also identified by blind docking experiments at the level of the membrane inner leaflet. Class II mutants discussed above (L69H, E56K, P67L, L206W and W361R) are also located in close vicinity or within this pocket, participating in its architecture and stability. Of note is that MD simulations indicate that L206W established strong interaction with P205 and W202, thereby moving the side chain of this last amino acid away from its position within the pocket in the wild-type situation, with W361 and W202 interacting with a same lipid (Fig. S11). Such modified interactions with lipids, as also likely for W361R (*18*), might account for a modified architecture of the pocket, that can be rescued by displacement of lipids by the corrector. E56 in the lasso Lh2 plays a role in the architecture/stability of the pocket through a salt bridge established with R75 in the elbow helix, forming the cytoplasmic side of the pocket. The 2,2-difluoro-1,3-benzodioxolyl portion of VX-809/VX-661 is deeply embedded in the pocket (Fig. S1), especially interacting with W361, S364, F81, F191 and M152, while the other portion of the drugs is oriented towards membrane lipids. The 2,2-difluoro-1,3-benzodioxolyl portion is common to VX-809, VX-661, as well as to the VX-809 derivatives ALK-809 and SUL-809, which shares a common mechanism of action (*40*). Of note is that among MSD1 class II mutations, E92K only showed full rescue at high concentrations (*15*), suggestive of a reduced affinity of the corrector, while G85E appeared resistant to correction (*19*). In the first situation, the salt-bridge formed by E92 with K95, both amino acids located at the pore-facing side of TM1, likely contributes to stabilize this helix. The E92K mutation would thus lead to increase MSD1 instability. In the second situation (G85E, facing the potential VX-809 binding site), alterations in TM1 span integration in the membrane was shown to be defective (*41*), possibly leading to the production of a non-rescuable CFTR. This was not the case of V232D which is corrected by VX-809 (*42*) and was recently associated with mis-insertion of the TM3/4 hairpin within the membrane (*43*). Interestingly, the hydrophobic pocket at the vicinity of which V232D is located, well discussed by Loo and Clarke (*42*), corresponds to the VX-445 binding site in MSD1 highlighted here at the level of the membrane outer leaflet (Fig. 6). The two pockets located in MSD1 at the level of the inner and outer membrane leaflets share a common hydrophobic character, whose disruption by mutations may be one of the mechanism responsible for a weakened stability and protein misfolding. Worth noting is that MD simulations have proposed that lipids can be stably accommodated in the putative VX-809 binding site studied here (*18*), thereby suggesting that VX-809 may mimic such molecules for stabilizing these fragile areas.

In the context of full-length CFTR-F508del, correction levels by VX-809 are however more modest (*15*) with only limited clinical benefits even in combination with the VX-770 potentiator (*9*). This limitation has been associated with additional NBD1 instability specific for the F508del mutation, which alters interdomain interactions of NBD1 occurring in particular through the MSD2 ICL4 loop (*44*). NBD1 binders, which may correct the instability of the domain, have been intensively searched and have led to highlight the interest of molecules which bind different sites on NBD1, such as c407 (*45*), crotoxin (*46*), nanobodies (*47*), BIA (5-bromoindole-3-acetic acid) and the dual corrector-potentiator CFFT-001 (*48–50*). Intensive investigations have been performed to identify molecules with strong affinity at the level of the two binding sites occupied by CFFT-001 and BIA (*51*), but were not successful. On the other hand, direct binding of VX-809 to NBD1 was demonstrated more recently (*24*). NMR data suggest that VX-809 occupies a site similar to that targeted by CFFT-001, and that allosteric coupling occurs between this site and the NBD1:ICL4 interface (*24, 49, 52*), while the associated mechanisms remain to be deciphered. While NBD1 thermo-instability was still detectable in the presence of VX-809, enhanced association to MSD1 via ICL4 favored protein assembly and CFTR-F508del rescue. Our results are consistent with the CFFT-001 binding pocket, highlighting the importance of not only H620 but also of Y625 for the binding of VX-809. This pocket could also be targeted by the more hydrophilic VX-445 corrector, which was recently shown to also directly bind and stabilize NBD1 (*13*), as H620 is here detected as a key amino acid implicated in VX-445 induced rescue. The architecture of this pocket might however depend on the conformational state of the CFTR regulatory (R) domain, which is located immediately downstream NBD1 helix H9. It should be noted here that another binding site was recently proposed for VX-809 and aminoarylthiazole-VX-809 hybrid derivatives, next to the ATP-binding site, in a location similar to the second scoring pose highlighted here by blind docking on the F508del-NBD1 3D structure (*53*). Whether or not this binding site is also relevant for increasing NBD1 stability and CFTR rescue deserves further investigation.

The NBD1 VX-809-binding site was shown to be allosterically connected to the F508 region interacting with ICL4 (*52*). Coupling between the MSD1 VX-809 binding site and the F508 region through a tight network involving amino acids from the elbow, lasso Lh2, ICL1 and ICL4 can be observed and supported here with secondary mutations. This network includes D47, E54, E56, R74, M152, K162, K163, K166 and E1075 (Fig. 6-left box). One key feature of allostery may be linked to the shift of the lasso Lh2 upon VX-809 binding and the associated interaction of D47 with K162, allowing a synergy between three communication pathways leading from MSD1 to K162 (the TM2 way: M152➔K162; two elbow-Lh2 ways: (i) R74➔R75➔ E56➔D47➔K162, (ii) R74➔R75➔E56➔E54➔K163➔K162) (Table S1). Such a network including ICL1 is also supported using truncation mutants (*54*). Recent experimental studies also highlighted the critical role that the lasso Lh2 is likely to play for CFTR assembly and pharmacological repair (*4, 21*). VX-809 binding onto the NBD1 site appears mandatory to rescue F508del in addition to occupancy of the MSD1 binding site. Also, class II mutants showed different sensitivity to the occupancy of these two sites. While F508del rescue can be abrogated by mutations in either one of the two sites, L206W rescue was affected by mutations only in the MSD1 site (Fig. 6).

It appears that VX-809 affects CFTR differently depending on the binding site, with the MSD1 site inducing a global stabilization of the initial folding steps while the NBD1 site stabilizes NBD1-ICL4 interaction. As VX-445 was shown to affect NBD1 folding (*13*) and binding of both VX-445 and VX-809 to MSD1 leads to increased allosteric coupling with NBD1, it can be suggested that both mechanisms contribute to the enhanced rescue observed with the VX-445/VX-809 combination.

## Acknowledgements

This work was performed using HPC resources from GENCI-[CINES] (Grants 2017-A0020707206, 2018-A0040707206, 2019-A0060707206 and 2020-A0080707206) and benefited from the financial support of the French Association Vaincre la Mucoviscidose and Association pour l’Aide à la Recherche contre la Mucoviscidose (AARM).

## Authors’ contributions

NB conceived the protocols, performed experiments and analysed data. AhE conceived the protocols and performed MD simulations, as well as data curation and analysis. JPM was involved in the conceptualization, design of the study and data analysis. BC, IP and NS performed experiments and statistical analysis. RZ and JLD participated in the analysis of the protein-drug interactions. AlE and ISG participated in the writing of the manuscript. IC and AH conceived and coordinated the project, conceived the protocols, performed experiments, analysed data and wrote the manuscript. All authors have reviewed the manuscript and approved its submission.

## Material and methods

### Plasmids and mutagenesis

The cDNA of CFTR WT (M470) was subcloned in pTracer as previously published (*55*). Mutagenesis of all the indicated mutants was performed using the QuickChange XL II mutagenesis kit (Agilent) following the manufacturer’s instructions. Obtained mutants were fully sequenced, amplified and purified (Macherey-Nagel). Plasmid concentrations were measured using a Nanodrop (Thermo Fisher Scientific).

### Cell culture and transfection

HEK293 cells were purchased from ATCC and cultivated in DMEM medium supplemented with 10% fetal calf serum (Thermo Fisher Scientific). Cells were maintained at 37°C, 5% CO2. For Western blot analysis, cells were seeded in 6-well plates and transfected with CFTR plasmids using Lipofectamine 3000 (Thermo Fisher Scientific).

### Western blot analysis

Transfected cells were treated with 3μM of VX-809 (Selleckchem) for 24 hours before being lysed in RIPA buffer containing protease inhibitors and protein concentration assessed using RcDc assay (BioRad). In some experiments cells were treated with 2μM of VX-445 (Selleckchem), alone and in combination with VX-809. Western blot analysis was performed using 60μg of protein from each sample separated on a 7% acrylamide gel. After transfer onto nitrocellulose membranes, CFTR was probed using antibody 660 (NACF Foundation) and alpha-tubulin probed with antibody DM1A (Santa Cruz).

### Statistical analysis

Quantitative variables were described as mean (± SEM). Comparisons to WT conditions and between treated and untreated conditions were made with one-way ANOVA followed by Fischer test for p evaluation.

### Visualization of 3D structures, molecular docking and molecular dynamics simulations

3D structures were visualized with the UCSF Chimera package (*56*). Formula of the drugs (Fig. S1) were extracted from the ZINC database (*57*). Blind docking was performed using the EADock DSS tool provided by the online SwissDock server (*30, 31*).

For molecular dynamics (MD) simulations, the 3D structure of human CFTR (pdb:6MSM) was embedded in a 1-palmytoyl-2-oleoyl-phosphatidylcholine (POPC) bilayer and solvated in a 150 mM NaCl solution. The CHARMM36 force field (*58*) was used for the protein, lipids and ions, and the TIP3P model for water. Minimization, equilibration and production steps were performed on the occigen/cines supercomputer using Gromacs 2019.1 (*59*). The standard CHARMM-GUI inputs (*60*) were used for the minimization and equilibration of the systems. During these steps, harmonic restraints applied to the protein heavy atoms and the lipid heads and were gradually released during 1.2 ns. The production dynamics were then performed in the NPT ensemble without any restraints. Nose-Hoover thermostat (*61*) and Parrinello-Rahman barostat (*62*) were used to keep the temperature and the pressure constant at 310 K and 1 bar. Periodic boundary conditions were used and the particle mesh ewald algorithm was applied to treat long range electrostatic interactions (*63*). A switching function was applied between 10 and 12 Å for the non-bonded interactions. LINCS (*64*) was applied to constrain the bond lengths involving hydrogen atoms. The integration timestep was set to 2 fs and the overall length of the trajectory was 1 μs for the simulation of apo CFTR.

MD simulations were also performed of the CFTR protein in complex with the correctors (VX-809/VX-661 and VX-445 ((4S)-methyl enantiomer in the trimethylpyrrolidin ring, as this one exhibits better efficacy in presence of VX-661 (*13*)) in the proposed binding sites, in order to evaluate the stability of the predicted conformations. Corrector charmm parameters were generated using the CgenFF module in CHARMM-GUI server (*65*). These MD simulations were run for at least 125 ns, with calculations of Root Mean Square Deviations (RMSD) done for estimating corrector stability within the binding sites. Binding poses at 125 ns were selected for illustration in the different CFTR-drug complexes.

The creation and analysis of CFTR networks (Fig. S6) were performed using the R package Bio3D (*66, 67*). In this method, MD simulations are used as an input that first undergoes a correlation analysis. Hence, a matrix of residue pairs dynamical cross-correlations is first calculated and the output is then used to generate a correlation network with residues as nodes that are linked by weighted edges that are proportional to their degree of correlated motion. The Girvan-Newman clustering method (*68*) is finally used to identify communities of highly correlated residues. The full residue network and coarse-grained community network were visualized and analyzed with VMD.

