## Supplemental Data for "CFTR corrector efficacy is associated with occupancy of distinct binding sites"

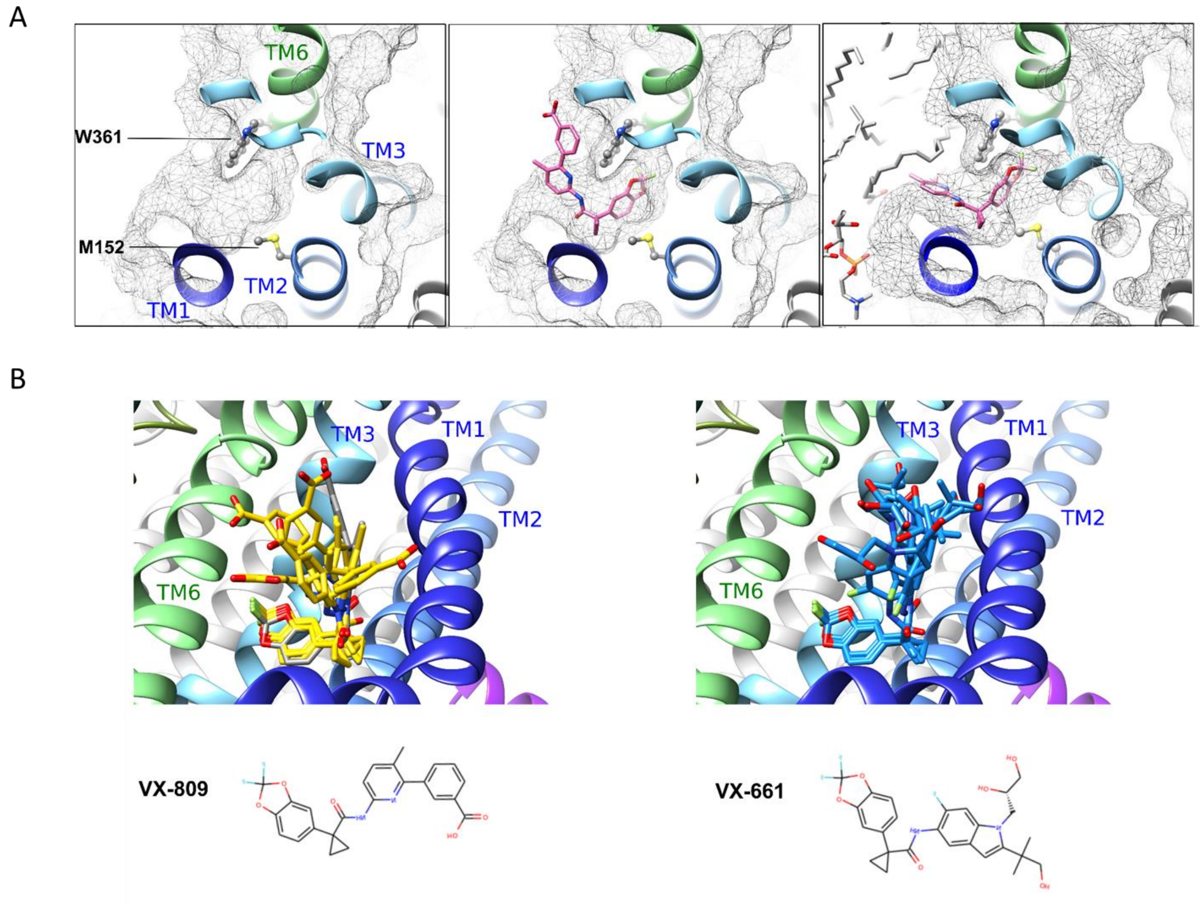


**Figure S1: Identification of a possible binding site for VX-809/VX-661 at the MSD1 inner membrane leaflet.**
**A.** View of the surface of the pocket at the MSD1 inner membrane leaflet, on the 3D structure of human CFTR (pdb 6MSM) - from left to right - before MD simulation, without and with VX-809 (best scoring pose using SwissDock) and after 125 ns MD simulation, embedded in lipids (POPC).
**B.** Main conformations of VX-809/VX-661 within the binding site at the MSD1 inner leaflet. Are shown at left the six most favorable VX-809 clusters, as identified using SwissDock. The structure of the first cluster of conformations is highlighted in grey, the others in yellow. These main conformations are compared to those observed (at right in blue) in the five most favorable clusters found for the docking of VX-661 on the same MSD1 3D structure, using SwissDock. *SwissDock FullFitness scores for these VX-661 clusters range between 1883 and -1880.*

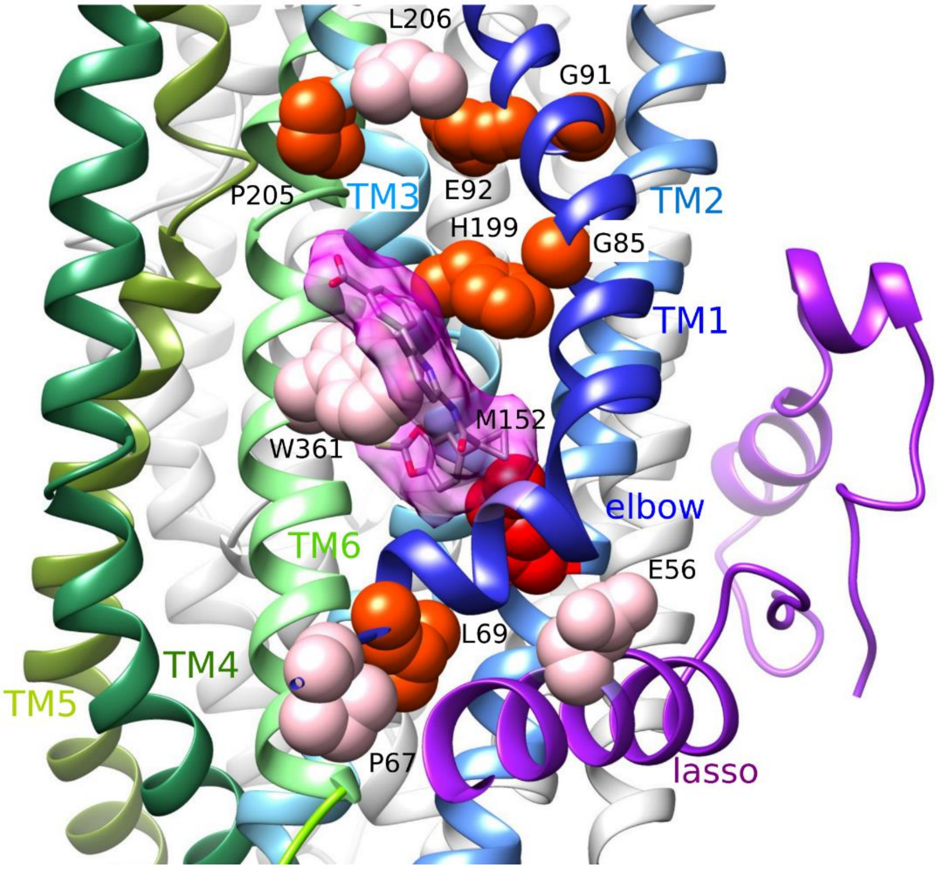


**Figure S2: Class II mutations in direct vicinity of the MSD1 VX-809 binding site**
Class II mutations are highlighted on the cryo-EM 3D structure of the full-length CFTR (pdb: 6MSM), with mild and severe mutations depicted in pink and orange, respectively.

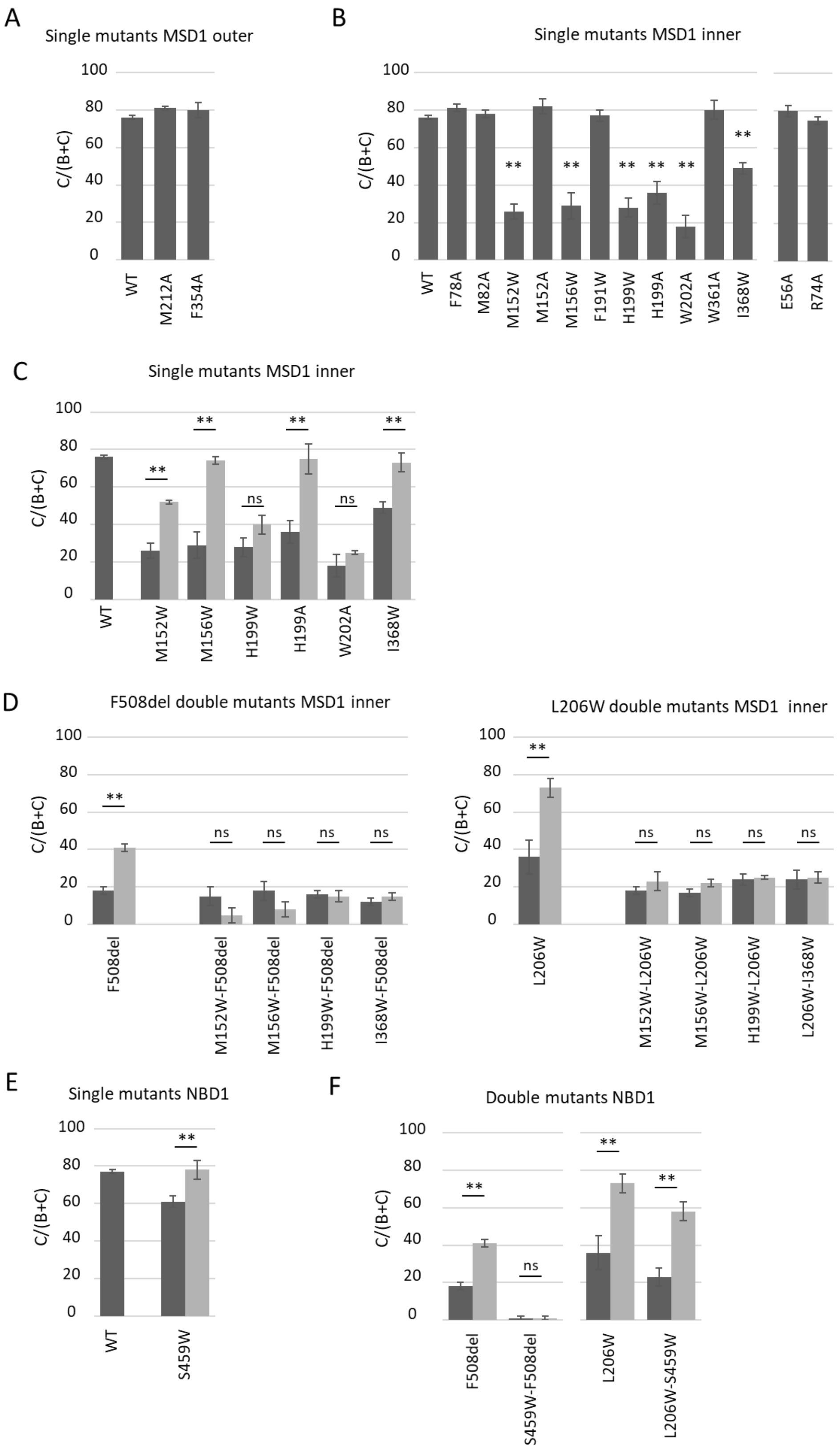


**Figure S3: Quantification of the C/(B+C) maturation ratio of the indicated mutants.**

**A-F.** Dark grey represents control conditions and light grey VX-809 treatments. Measures are means ± SEM of n=3-10, with * indicating $p<0.05$, ** indicating $p<0.01$ and ns indicating no statistical differences (one-way ANOVA followed by Fischer test for p evaluation).

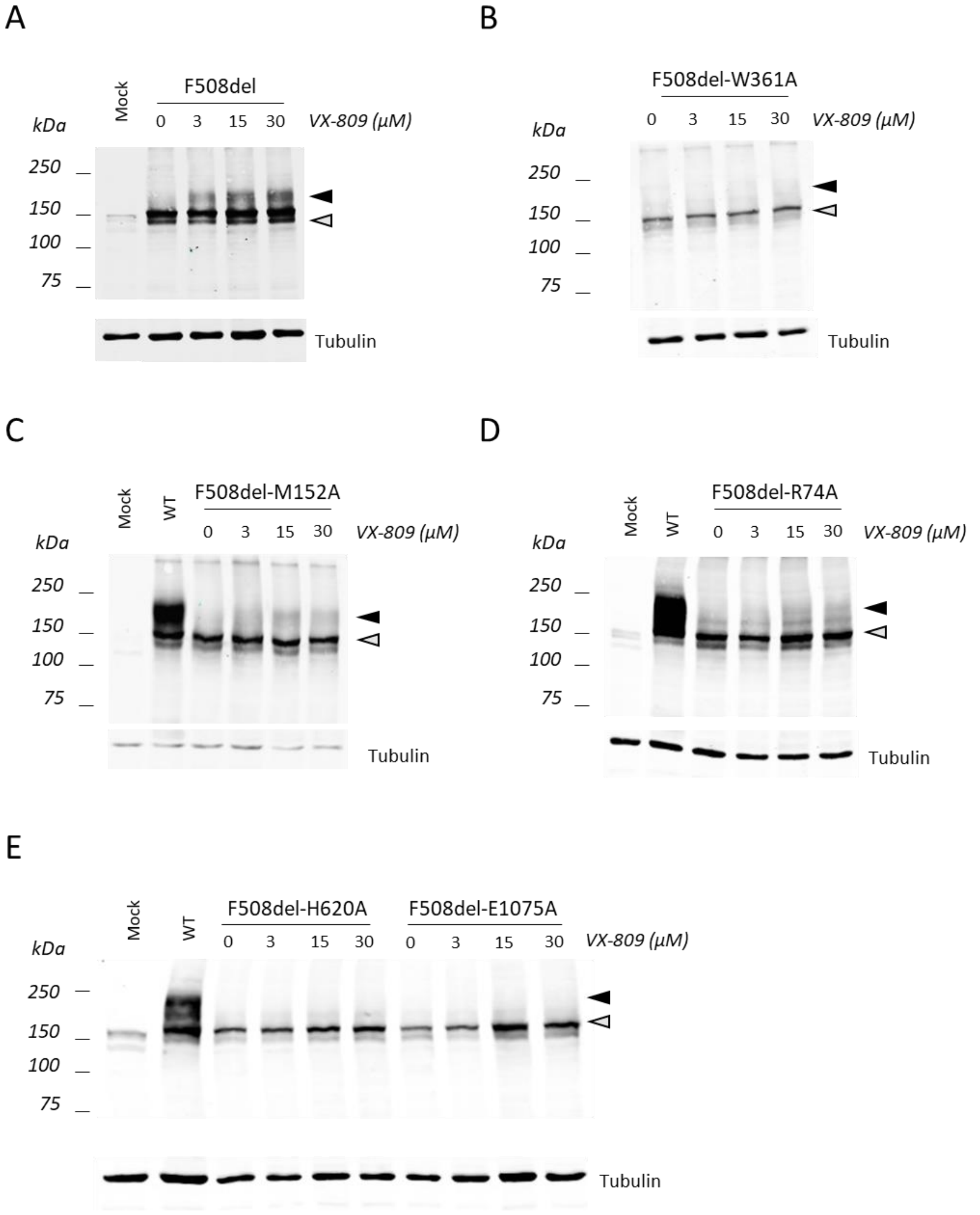


**Figure S4: Western blot analysis of F508del and F508del double mutants.**
**A-E.** When indicated, cells were treated with increasing concentrations of VX-809 (3, 15 or 30 µM, 24h). Grey arrow indicates core glycosylated CFTR (band B) and black arrow fully glycosylated CFTR (band C). Alpha-tubulin was probed to assess equal protein loading.

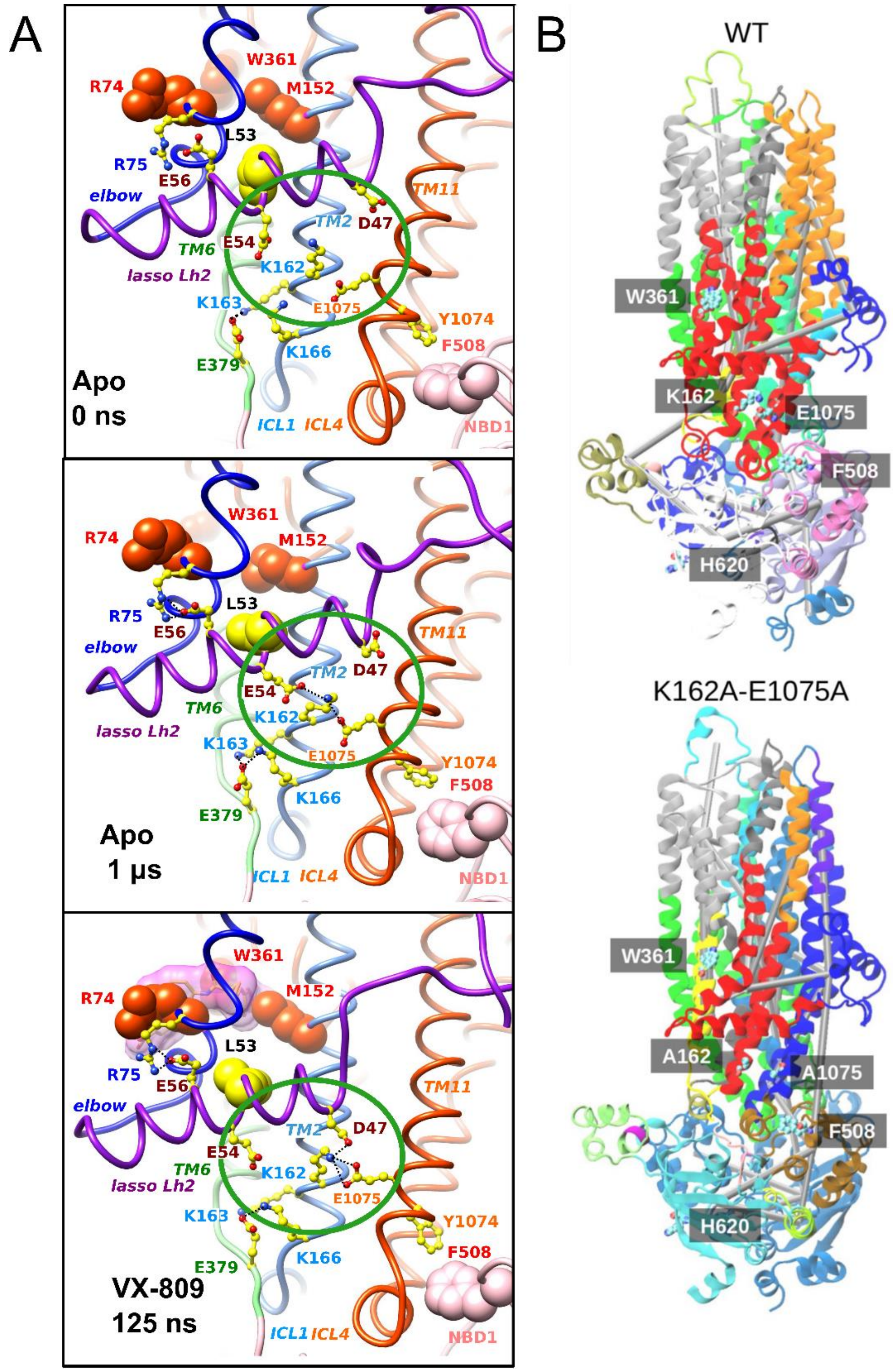


**Figure S5 : Allosteric coupling between MSD1 and NBD1.**

**A.** Focus on the lasso Lh2-ICL1-ICL4 network. From top to bottom: 3D structure of the human CFTR 3D structure in absence of VX-809, before (0 ns) and after (1 µs) MD simulation, and in presence of VX-809 in the inner membrane leaflet site (125 ns MD simulation). A shift (~5 Å) of lasso Lh2 relative to ICL1/ICL4 is observed in the presence of VX-809, allowing K162 to bind both D47 and E1075. This shift, which appeared rapidly in the beginning of the MD simulation and stayed very stable, may be a critical feature of the allosteric communication between the MSD1 binding site and ICL4.

**B.** Dynamical network analysis of the wild-type (top) and double mutant K162A-E1075A (bottom) protein 3D structures (1 μs MD simulation), highlighting communities of highly correlated residues. Each community is represented by one color, with cylinders connecting the critical nodes. The community highlighted in red in the wild-type protein, connecting the VX-809 binding site on MSD1, including W361, to ICL4, in contact with NBD1 F508, is broken in the mutated one (red and blue colors).

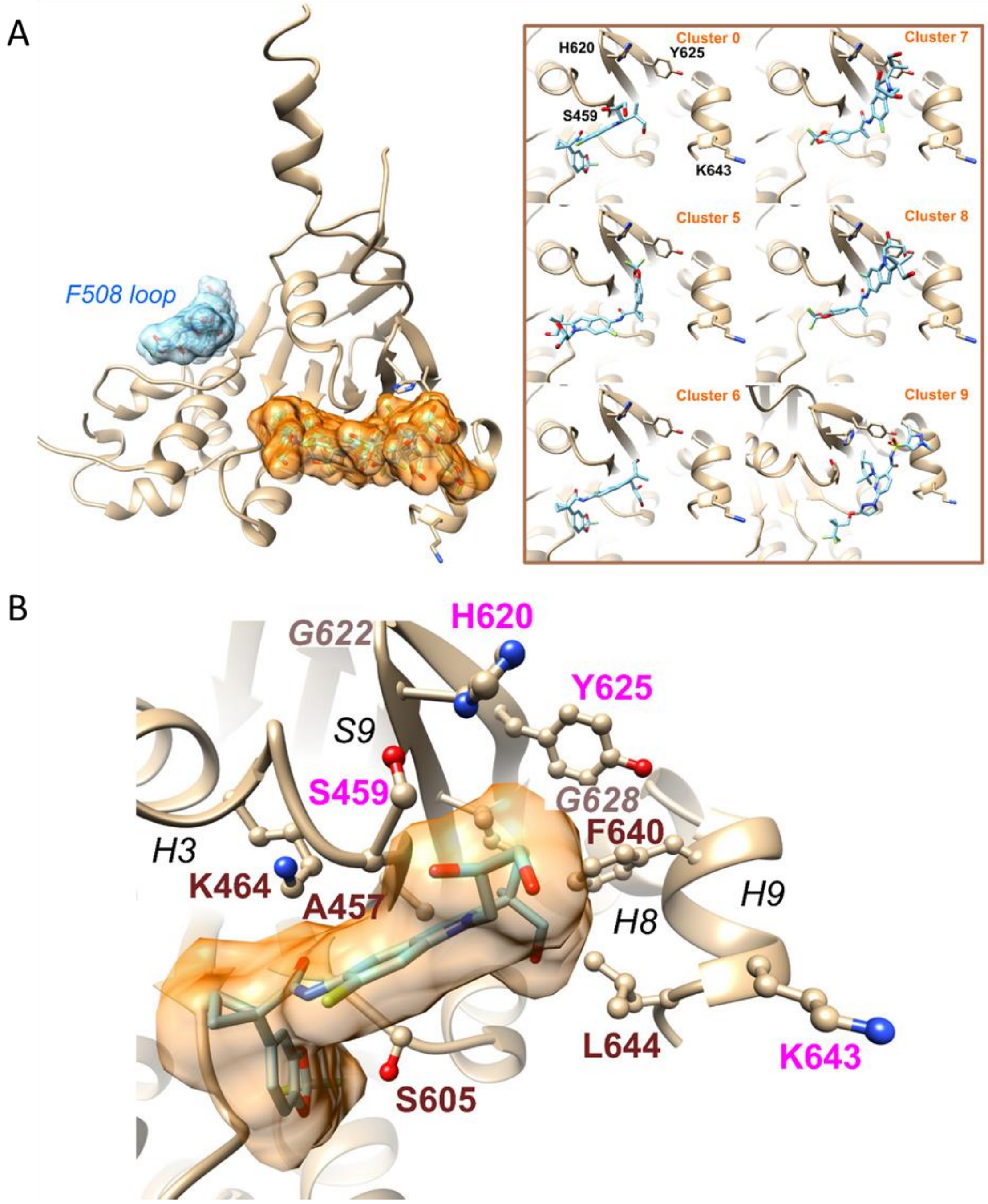


**Figure S6: VX-661 binding site in NBD1**

**A.** View of the human CFTR F508del NBD1, extracted from a crystal structure (pdb: 4WZ6) from which the regulatory extension (RE) has been removed and on which VX-661 has been docked (blind docking). Two main pockets were here identified in the ten high-ranking clusters, the second one (orange) gathering 6 clusters out of 10 *(SwissDock FullFitness scores of -1224 to 1217)*. Representative conformations of each of the 6 clusters, with the main amino acids of the binding pocket highlighted in ball and stick. The conformation of the cluster with the best docking score is detailed at bottom **(B)**.

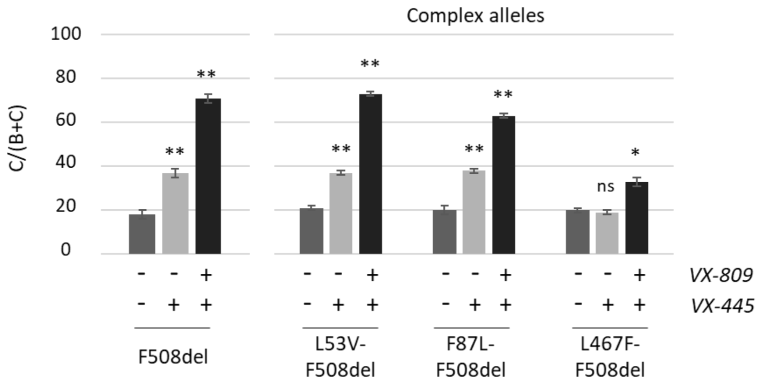


**Figure S7: Quantification of the C/(B+C) maturation ratio of the indicated mutants.** Dark grey represents control conditions and light grey VX-809 treatments. Measures are means ± SEM of n=3-10, with * indicating p<0.05, ** indicating p<0.01 and ns indicating no statistical differences (one-way ANOVA followed by Fischer test for p evaluation).

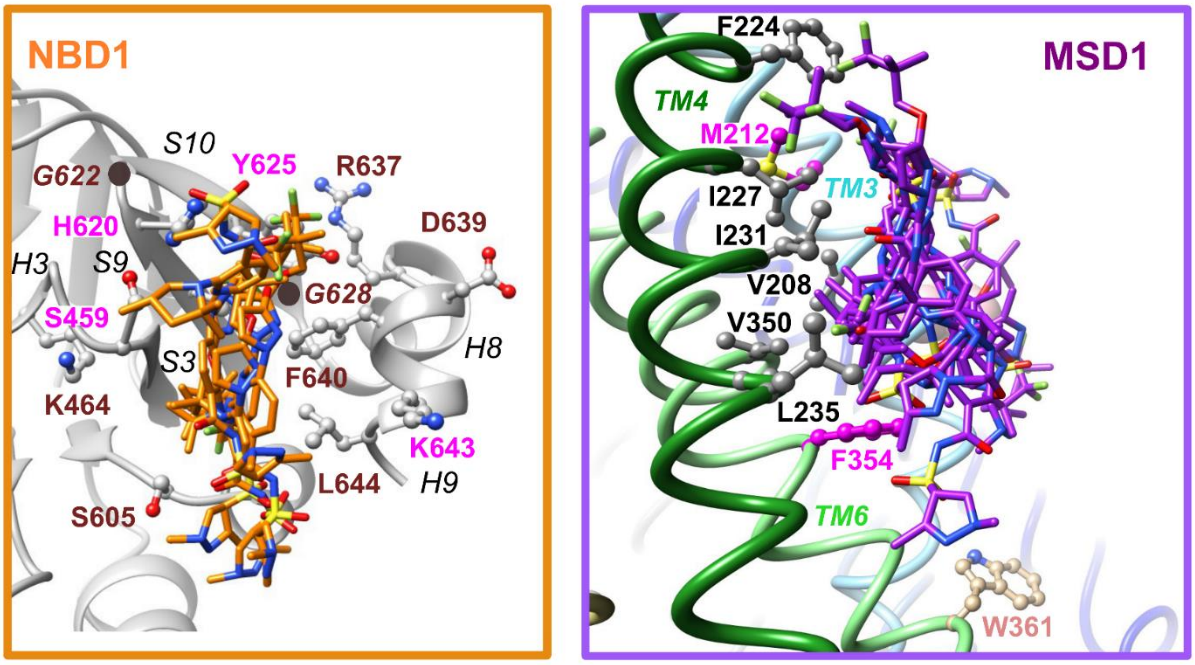


**Figure S8:** Representative conformations of the SwissDock clusters with the best scores in MSD1 and in NBD1, and amino acids participating in the pockets.

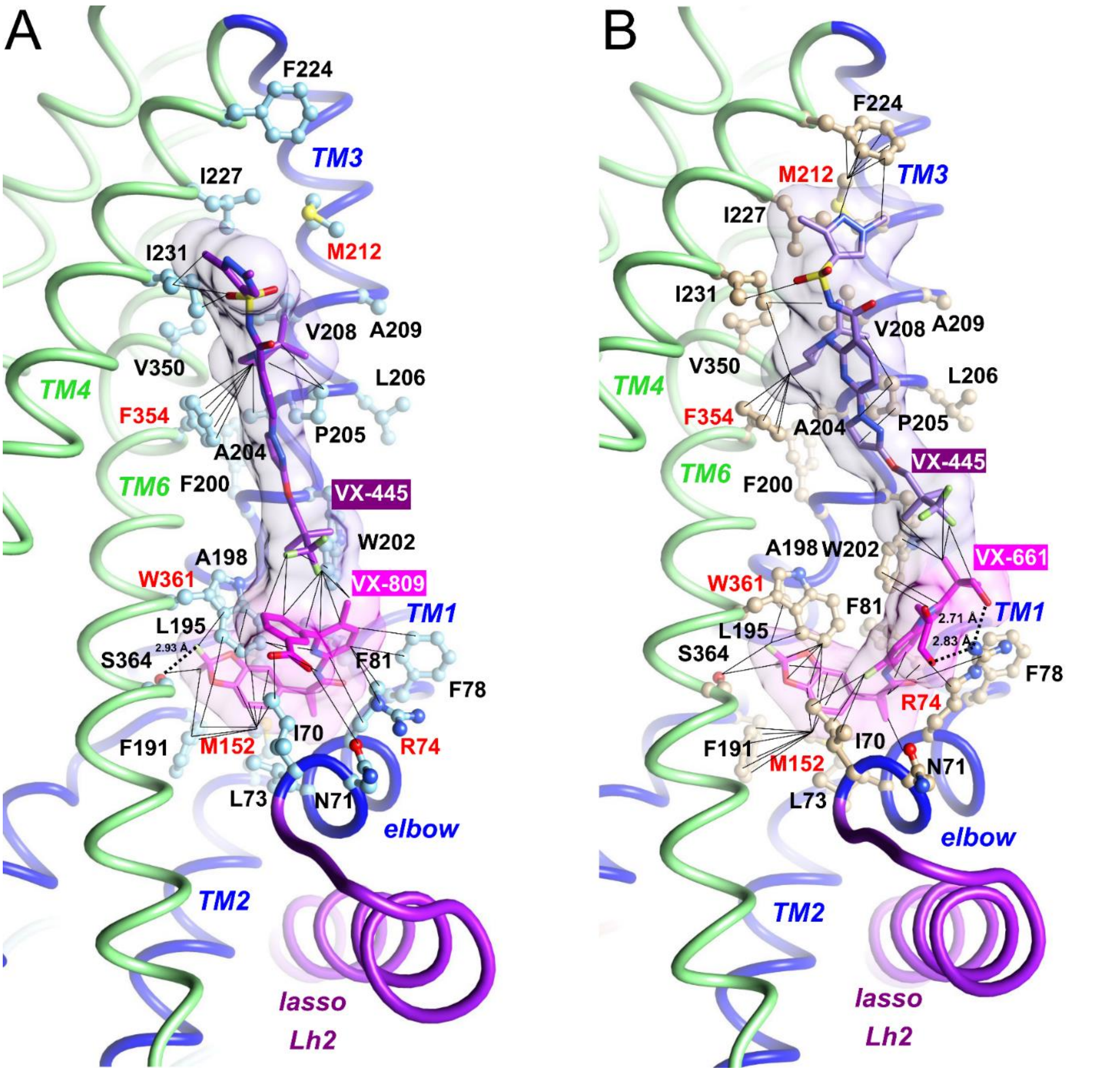


**Figure S9: VX-445 and VX-809/VX-661 in MSD1**

**A.** View after 125 ns MD simulation of the VX-445:VX-809 complex within human CFTR MSD1. Main contacts are visualized by black tiny lines. Of note is the interaction of the 3,3,3-trifluoro-2,2-dimethylpropoxyl group a one extremity of VX-445 with VX-809, as well as the interaction of (4S)-methyl of the VX-445 trimethylpyrrolidin ring with F354. A H-bond is likely to occur between VX-809 and S364 (2.93 Å).

**B.** View after 125 ns MD simulation of the VX-445:VX-661 complex within human CFTR MSD1. R74 makes here two H-bonds with VX-661 (2.71 Å and 2.83 Å). The extremity of VX-661 also links the 3,3,3-trifluoro-2,2-dimethylpropoxyl group at one extremity of VX-445, whereas the other extremity of VX-445 strongly interacts with F224, which is stably positioned through a strong interaction with M212.

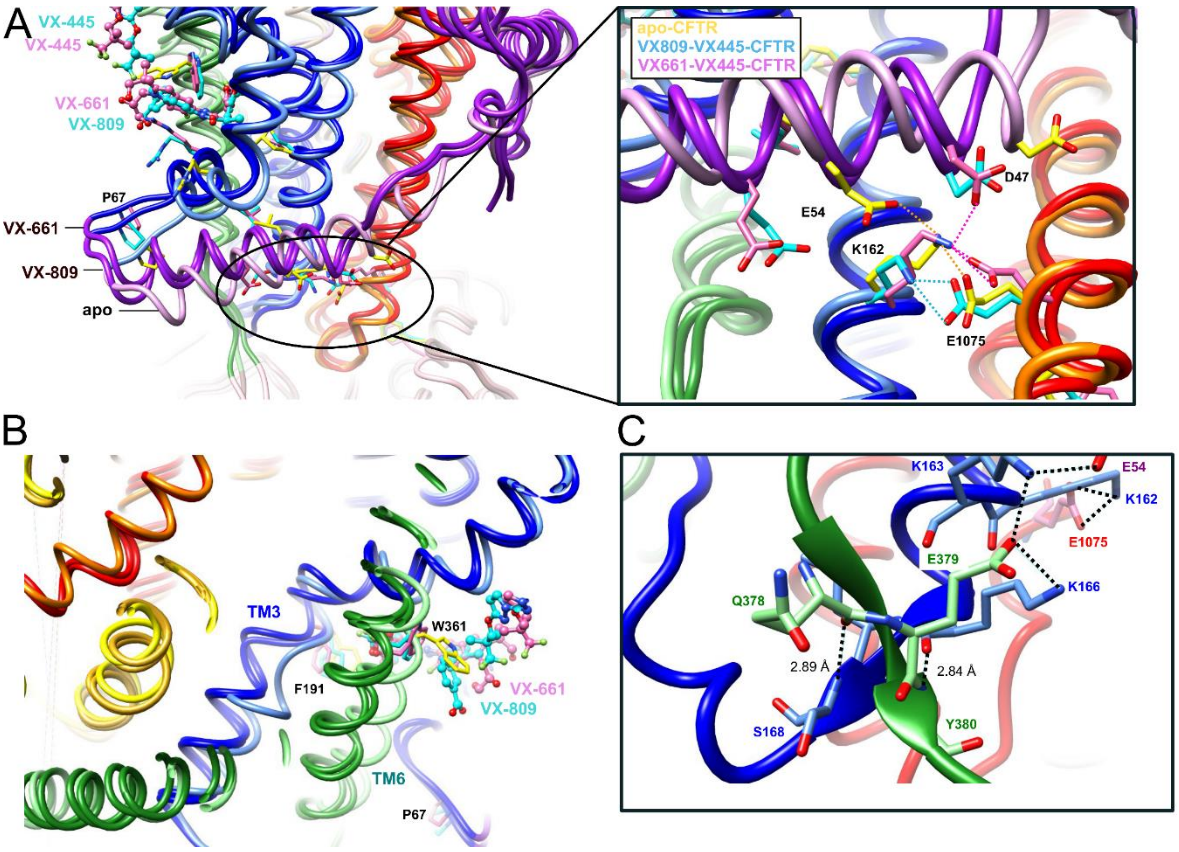


**Figure S10: Behavior of the lasso Lh2-ICL1-ICL4 region in the presence of two correctors.**
**A.** Superimposition of the human CFTR 3D structures in complex with VX-661/VX-445 and VX-809/VX-445 (after 125 ns MD simulation) and without corrector (apo, 1 μs MD simulation, ribbons colored lighter). The conformation of apo CFTR after 1 μs MD simulation is similar to that observed after 125 ns, and remains stable over time. The view illustrates the movements occurring in the vicinity of VX-809/VX-661, in particular at the level of the elbow helix and the lasso Lh2. A shift of ~5 Å is observed for Lh2, allowing D47 to bind K162, which binds E1075.
**B.** Movements occurring in the region of TM3-TM6, which includes F191 and W361.
**C.** Focus on the region including K166, in the VX-661/VX-445 CFTR complex, showing a small, cross β-sheet formed between K166-S168 (ICL1) and Q378-Y380 (end of TM6). This striking feature is likely the consequence of a particularly good conformation of MSD1 with this particular combination of the two correctors.

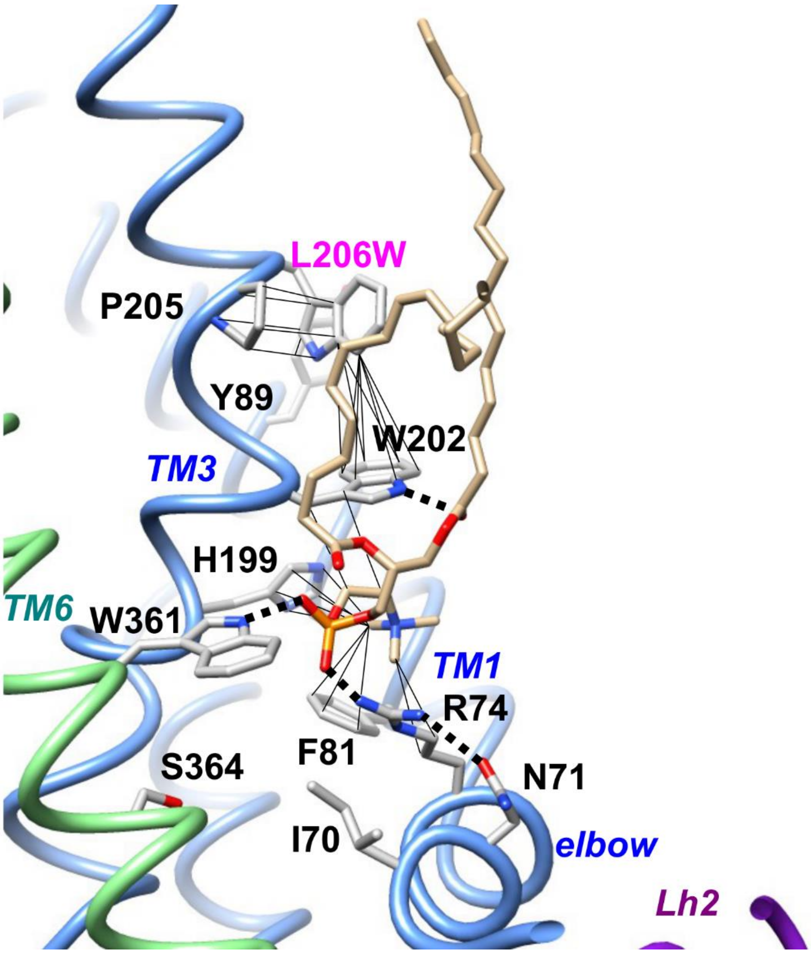


**Figure S11: Possible impact of the L206W mutation, as appreciated after short MD simulation (125 ns)**.

MD simulation led to a rapid stabilization of amino acids within the inner membrane pocket. L206W, one helix turn from W202, is directly followed by P205, which orientates, through stacking, the side chain of L206W. This leads to also attract W202 side chain, which leaves its position of the wild-type situation (rotamer in contact with W361). Hence, it leaves room for the polar head groups of a lipid molecule (POPC), which is stabilized by multiple contacts with R74, F81, H199, W202 and W361. Once the corrector occupies the pocket, this lipid is likely to be displaced and W202 able to adopt a rotamer side chain position similar to that of the wild-type CFTR.

| | ***VX-661+VX-445*** | ***VX-809*** | ***VX-809+VX-445*** | ***VX-661*** | ***VX-445*** |
|---|---|---|---|---|---|
| *ICL1 K162* | ++ | ++ | ++ | + | + |
| *Lasso Lh2 D47* | ++ | ++ | + | - | - |
| *Lasso Lh2 E54* | ++ | + | + | + | - |
| *Cross β-sheet* | ++ | ++ | - | ++ | ++ |

**Table S1: Allosteric linkage observed after MD simulation (125 ns) of human CFTR in presence of corrector(s) in the MSD1 binding site(s).** This linkage, with reference to central amino acids of the network, as depicted in Fig. 3, was appreciated based on the presence (++/+) or absence (-) of H-bonds and/or salt-bridges allowing direct communication. ++ and + distinguish bonds which are well formed and those allowed by one standard side chain rotamer, respectively. The cross β-sheet refers to the architecture depicted in Fig. S9C, between the end of TM6 and ICL1.
